# A newly identified group of P-like (PL) fimbriae from extra-intestinal pathogenic *Escherichia coli* (ExPEC) encode distinct adhesin subunits and mediate adherence to host cells

**DOI:** 10.1101/2021.07.20.453171

**Authors:** Hajer Habouria, Hicham Bessaiah, Julie Buron, Sébastien Houle, Charles M. Dozois

**Affiliations:** INRS-Centre Armand-Frappier Santé Biotechnologie, Laval, Québec, Canada; CRIPA-Centre de recherche en infectiologie porcine et avicole, Saint-Hyacinthe, Québec, Canada

**Keywords:** fimbriae, pili, pathogenic *E. coli*, hemagglutination, biofilm, adherence, urinary tract infection, poultry

## Abstract

Fimbrial adhesins play a critical role for bacterial adherence and biofilm formation. Sequencing of avian pathogenic *Escherichia coli* (APEC) strain QT598 identified a fimbrial gene cluster belonging to the π group that we named PL (P-like) fimbriae, since genetic organization and sequence are similar to Pap and related fimbriae. Screening of genomic databases indicated that genes encoding PL fimbriae located on IncF plasmids are present in a diversity of *E. coli* isolates from poultry, human systemic infections, and other sources. As with P fimbriae, PL fimbriae exhibit sequence divergence in adhesin encoding genes, and strains could be divided into 5 classes based on differences in sequences of the PlfG adhesin protein. The *plf* genes from two predominant PlfG adhesin classes, PlfG-I and PlfG-II were cloned. PL fimbriae were visualized by electron microscopy, promoted biofilm formation, demonstrated distinct hemagglutination profiles and promoted adherence to human bladder and kidney epithelial cell lines. Hybrid fimbriae comprised of genes from *plf_QT598_* wherein *plfG* was replaced by *papG* encoding adhesin genes were also shown to be functional and mediate adherence to epithelial cells, further indicating similarity and functional compatibility between these two types of fimbriae. Although deletion of *plf* genes did not significantly reduce colonization of the mouse urinary tract, *plf* gene expression was increased over 40-fold in the bladder compared to during in vitro culture. Overall, PL fimbriae represent a new group of fimbriae demonstrating both functional differences and similarities to P fimbriae and may contribute to adherence to cells and colonization of host tissues.

**Importance:** Fimbriae are important colonization factors in many bacterial species. The identification of a new type of fimbriae encoded on some IncF plasmids in *E. coli* was investigated. Genomic sequences demonstrated these fimbrial gene clusters have genetic diversity, particularly in the adhesin encoding PlfG gene. Functional studies demonstrated differences in hemagglutination specificity, although both types of Plf adhesin under study mediated adherence to human urinary epithelial cells. Such fimbriae may represent previously unrecognized adhesins that could contribute to host specificity and tissue tropism of some *E. coli* strains.

## Introduction

Bacterial adherence to surfaces is an important survival mechanism. Attachment to host cells or extracellular matrix can provide access to specific niches and promote colonization of host tissues. Adhesins can also mediate biofilm formation through bacteria-bacteria associations and improve survival in the environment. Bacterial adhesins include hair-like appendages (fimbriae or pili) as well as other molecules, including proteins or polysaccharides, which are displayed on the cell surface (Soto *et al*., 1999). Many types of fimbriae (pili) in Gram-negative bacteria are assembled by the chaperone/usher pathway (CUP) (Nuccio *et al*., 2007, Werneburg *et al*., 2018). In *Escherichia coli*, numerous types of CUP fimbriae have been identified and characterized. Pathogenic *E. coli* often can produce multiple types of fimbrial adhesins, and genomic analyses indicate that some strains may contain 10 or more fimbrial gene clusters (Werneburg *et al*., 2018, Wurpel *et al*., 2013). The ability to produce a variety of adhesins can provide a fitness advantage by expanding potential host receptor targets, or promoting adherence to environmental substrates.

Two main types of fimbriae, type 1 and P fimbriae, from *E. coli* have been extensively investigated to determine aspects of their roles in disease, particularly urinary tract infections (Ambite *et al*., 2019, Connell *et al*., 1996, Sokurenko *et al*., 1992), as well as molecular aspects of biogenesis and assembly of these structures (Lillington *et al*., 2014). Type 1 fimbriae mediate adherence to mannose-containing receptors, and have been shown to be critical for virulence of extraintestinal pathogenic *E. coli* (ExPEC) including *E. coli* strains causing urinary tract infections (Gunther *et al*., 2002), and neonatal meningitis (Khan *et al*., 2007). P fimbriae were first described in uropathogenic *E. coli* (UPEC) and were named based on receptor affinities for P blood group oligosaccharides, and were also described as Pyelonephritis-associated pili (Pap), since these fimbriae were more associated with *E. coli* strains from cases of pyelonephritis (Plos *et al*., 1990). P fimbriae have also been identified in other ExPEC including *E. coli* associated with systemic infections in swine (Dezfulian *et al*., 2003) and some strains of avian pathogenic *E. coli* (APEC) (Dozois *et al*., 1995, Kariyawasam *et al*., 2006, Mellata *et al*., 2003, van den Bosch *et al*., 1993). The P fimbrial gene clusters are commonly located on horizontally-acquired chromosomal regions, that have been termed pathogenicity islands (Blum *et al*., 1995, Guyer *et al*., 1998, Kariyawasam *et al*., 2006).

The P fimbrial gene cluster comprises 11 genes including regulatory genes (*papI* and *papB*) and genes dedicated to fimbrial assembly and structure (*papAHCDKJEFG*). The *papA* gene encodes the major fimbrial subunit, and has been used to class P fimbriae into serological variants (F7_1_, F7_2_ through F16) (Johnson *et al*., 2000). The adhesin-specificity of P-fimbriae is mediated by the *papG* gene product. The G adhesins of P fimbriae were grouped into 3 major classes based on sequence differences and receptor specificity to different Gal(α1-4)Gal-containing glycolipids (Marklund *et al*., 1992, Stromberg *et al*., 1990). PapG I recognize globotriaosylceramide or GbO3, PapGII recognize globotetraosylceramide (GbO4) and PapGIII or PrsGIII recognize galactosylgloboside (GbO5). These glycolipid receptors are usually found on the surface of red blood cells and on human bladder and kidney cells. A distinct variant allele, *papG*_BF31_, which was termed as class IV was also reported, although receptor specificity for this fimbrial adhesin was not described (Manning *et al*., 2001).

P fimbriae are the archetype representatives of the π fimbrial family (Nuccio *et al*., 2007), that includes a number of other types of *E. coli* fimbriae including Pix fimbriae present in some UPEC strains (Lugering *et al*., 2003, Schneider *et al*., 2004) and Sfp fimbriae encoded on plasmids in some lineages of enterohemorrhagic *E. coli* (Bielaszewska *et al*., 2009, Brunder *et al*., 2001). This report describes a new type of *E. coli* fimbriae from the π group that we have named P-like (PL) fimbriae, since they share sequence similarity and genetic organization with P fimbriae. The PL fimbriae are distinct from other known members of the π fimbriae and are encoded on IncFIB plasmids containing numerous other virulence genes associated with ExPEC and APEC strains. As with P fimbriae, PL fimbriae have also diversified into a number of different G adhesin classes and major subunit variants, suggesting adaptive potential for host specificity and tissue tropism. We characterized two different types of PL fimbriae encoding distinct G adhesins, and demonstrate these fimbriae can mediate adherence to host epithelial cells.

## Results

### Genomic analysis identifies a new type of fimbriae with a genetic organization similar to P fimbriae

Previously, we reported that ExPEC strain QT598, originally isolated from an infected turkey, contains a large ColV-type virulence plasmid, pEC598, that encodes a novel autotransporter protein, the **<u>s</u>**erine-protease **<u>h</u>**emagglutinin **<u>a</u>**utotransporter, Sha (Habouria *et al*., 2019). The region adjacent to the *sha* gene on pEC598 encodes a fimbrial gene cluster (Fig. 1). Due to the close protein identity and genetic organization with P fimbriae (see below), we have named these genes *plf* (for **<u>P</u>**-**l**ike **<u>f</u>**imbriae) and called these adhesins PL fimbriae. The *plf* gene cluster is inserted beside a site-specific integrase gene located adjacent to the REP FIB region on the pEC598 plasmid (Fig. 1). The REPFIB region and *intM* integrase genes are also present in most IncFII plasmids, and are also commonly flanked by other predicted virulence genes such as *mig14*, *hlyF*, and *ompT* in other APEC virulence plasmids, suggesting this conserved region may have led to acquisition of different genes through integration/recombination. Interestingly, this region of F and related plasmids is considered a « hot spot » for insertion of a diversity of virulence and antibiotic resistance genes, that have been termed « cargo genes » (Fig. 1) (Koraimann, 2018).

**Fig. 1.**
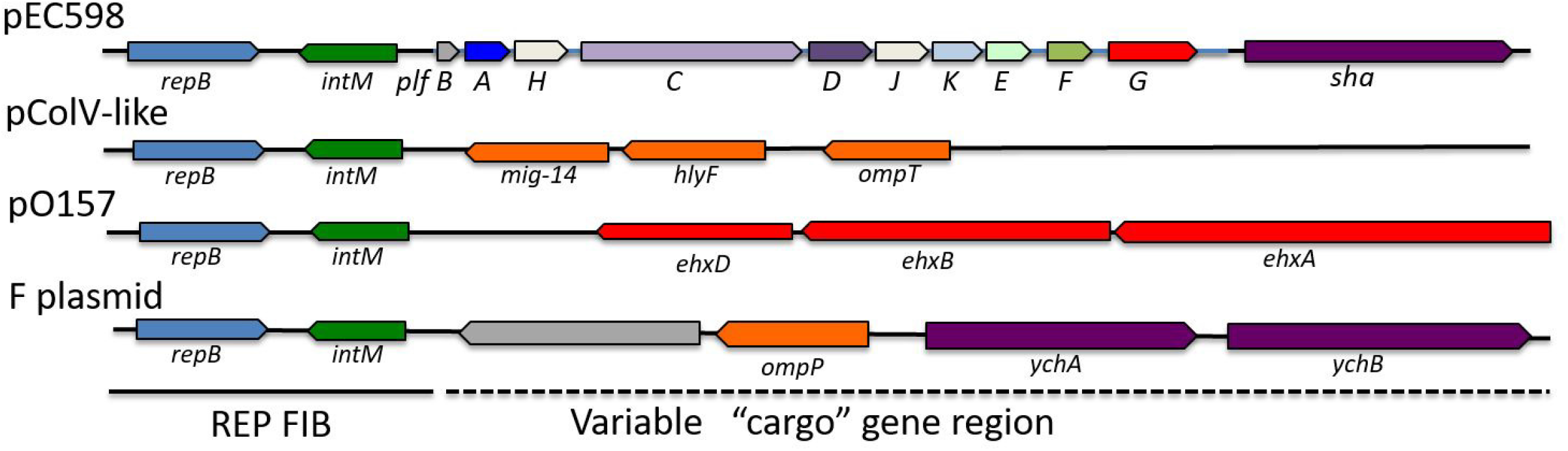
The P-like fimbrial (*plf*) gene cluster of plasmid pEC-598-1 is located adjacent to the REPFIB region. Comparison of the cargo genes adjacent to the REPIB region on F- and related conjugative plasmids. In other ColV-like plasmids, such as pAPEC-1 or pVM1, the cargo region encodes *mig-14*, *hlyF*, and *ompT* virulence genes. In pO157 from *E. coli* O157:H7 strains, the genes encoding the Ehx hemolysin RTX-toxin are within the cargo region. The F-plasmid cargo gene region also encodes virulence-associated genes, a protease, OmpP, that can degrade host defense peptides and genes encoding self-associating AIDA-1 like autotransporters, YchA and YchB. NCBI Accession numbers: pEC598 (NZ_KP119165.1), “pColV-like”: pAPEC-1 (CP000836), pVM01 (NC_010409.1), pO157 (AB011549), F plasmid (NC_002483.1).

Fimbriae have been classed into specific groups by a variety of criteria, including comparison of usher, chaperone, and major fimbrial subunits (Girardeau *et al*., 2000, Nuccio *et al*., 2007). Fimbriae classification using the usher encoding protein sequences has placed P fimbriae within the π fimbrial clade (Nuccio *et al*., 2007). Phylogenetic analysis using the usher proteins indicated that the PL fimbriae also belong to the π fimbrial clade and cluster with Pix, Sfp and Pap fimbriae (Fig. 2A). Phylogenetic comparison based on the chaperone proteins also indicated PL fimbriae were most closely related to P, Pix, and Sfp fimbriae (Supplemental Fig. S1).

**Fig. 2.**
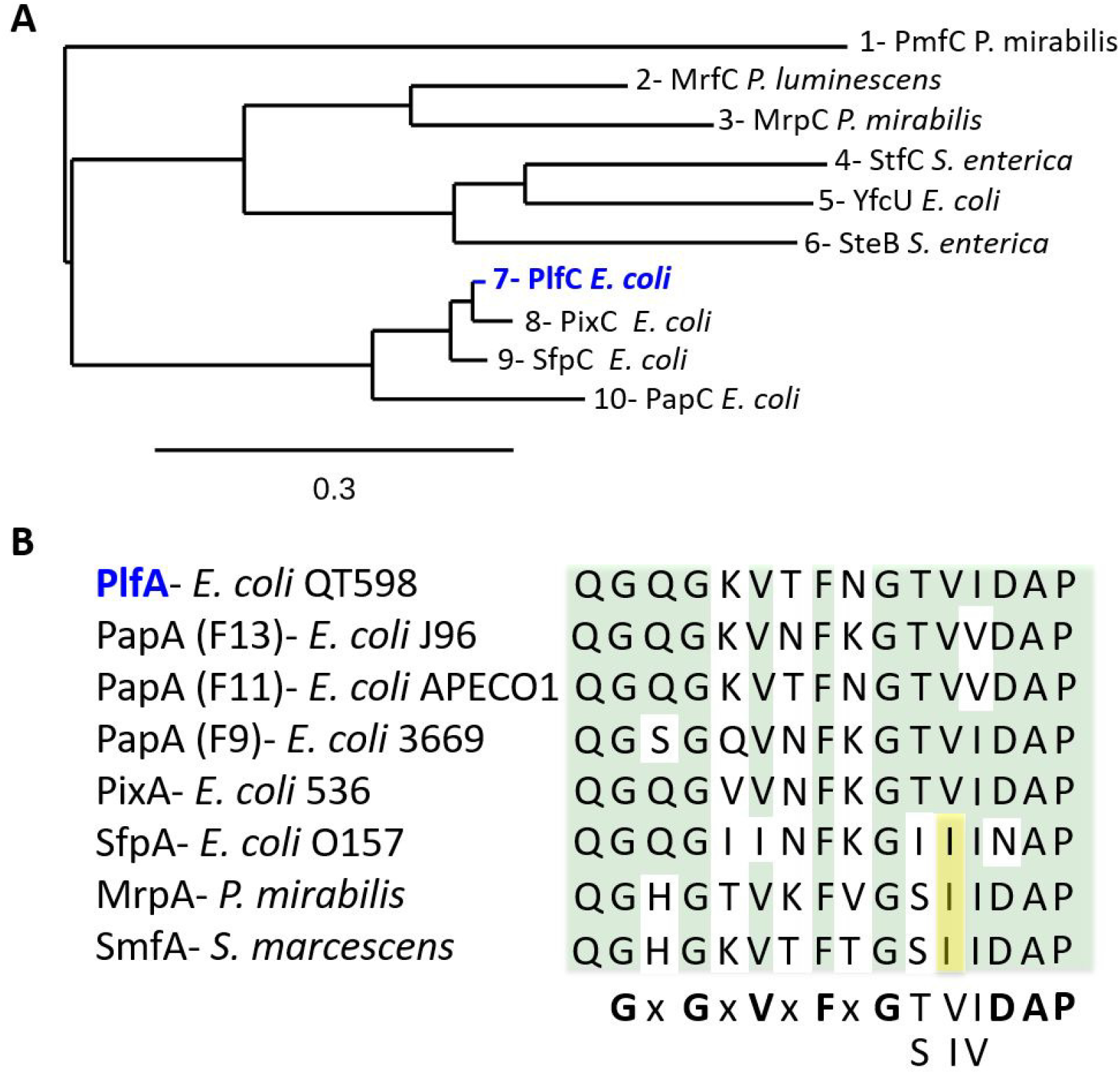
Phylogenetic relationship of PL fimbriae (Plf) with other types of fimbriae. A. Phylogram using sequences of fimbrial usher proteins (FUPs) belonging to the π fimbriae clade based on the classification scheme of (Nuccio *et al*., 2007). The PlfC protein, shown in blue, clusters with other FUPs, Sfp and Pix, more closely related to P fimbriae. **B.** Alignment of segment S1 of the major subunit proteins also places PlfA within the PapA-like subfamily (Ic) according to the scheme of Girardeau *et al*. (Girardeau *et al*., 2000). The alignment results in a consensus (**G**x**G**x**V**x**F**xG[TS][VI][IV] **DAP**) motif. Sequences used for **(A)** were UniProt (uniprot.org): **1-**P53514, **2-**Q93MT4, **3-**Q51904, **4-**H9L4A4, **5-**P77196, **6-**A0A3U8KZY6, **7-** Genbank-NCBI: AKG46878.1, **8-** A0A454A7L3, **9-** B8RHG0, **10-** P07110. Sequences used for **(B)** were **PlfA**-Genbank-NCBI: AKG46876.1; UniProt (uniprot.org) **PapA (F13)**- X61239, **PapA (F11)**-Q4FBG1, **PapA (F9)**-M68059, **PixA**-A0A454A7E1, **SfpA-** W6JHT1, **MrpA-** Q03011, **SmfA**-P13421.

Girardeau *et al*. (Girardeau *et al*., 2000) also classified fimbriae based on amino acid sequence motifs within the major subunit proteins. The subfamily Ic (PapA-like) group which included PapA variants, SmfA (*Serratia marscesens*), PmpA (*Proteus mirabilis*) and MrpA (*Proteus mirabilis*) also includes PixA, SfpA, and PlfA major subunit proteins that share a conserved sequence signature motif in segment S1 of the fimbrial subunits: **(GxG[KT]V[TS]FxG[TS]V[VI]DAP**) (Fig. 2B).

The PL fimbriae (*plf*) gene cluster contains 10 genes predicted to encode one regulatory and 9 structural/assembly proteins that share identity to equivalent proteins present in the *pap* gene cluster (Fig. 3). A predicted regulatory protein PlfB, shares identity to members of the PapB regulatory family that includes PapB (P fimbriae), PixB (Pix fimbriae), FocB/SfaB (F1C/S fimbriae), AfaA (Afa-III adhesin), Daa (F1845 fimbriae), and FanA/FanB (K99 pili) regulatory proteins (Holden *et al*., 2001). The highest identity was to PixB (57% identity / 76% similarity), followed by FanB (47% identity / 69% similarity) and PapB (45% identity / 67% similarity). No equivalent of the PapI regulatory gene was present. Some Plf proteins show higher identity to other *pap*-related fimbrial gene clusters, specifically from Pix fimbriae, identified in some *E. coli* urinary tract infection isolates (Lugering *et al*., 2003, Schneider *et al*., 2004), and the plasmid-encoded Sfp fimbriae, present in sorbitol–fermenting diarrheagenic *E. coli* O157:H7 strains (Bielaszewska *et al*., 2009, Brunder *et al*., 2001) (Fig. 3). Despite demonstrating higher identity to certain proteins from these other fimbriae, only Plf demonstrates a complete set of structural/assembly proteins equivalent to each in the *pap* gene cluster, as Pix and Sfp fimbriae both lack PapE or PapK protein paralogs, which are known to code for a fibrillum subunit and adaptor protein respectively (Fig. 3). The gene products showing the greatest degree of diversity among these fimbrial protein paralogs were the G adhesins that exhibited less than 30% identity. Taken together, the PL fimbrial system is highly similar to Pix and Sfp fimbrial systems, but shares a genetic organization more akin to P fimbriae, as it includes the PapK and PapE paralogous proteins PlfK and PlfE predicted to be part of a thin fibrillum structure.

**Fig. 3.**
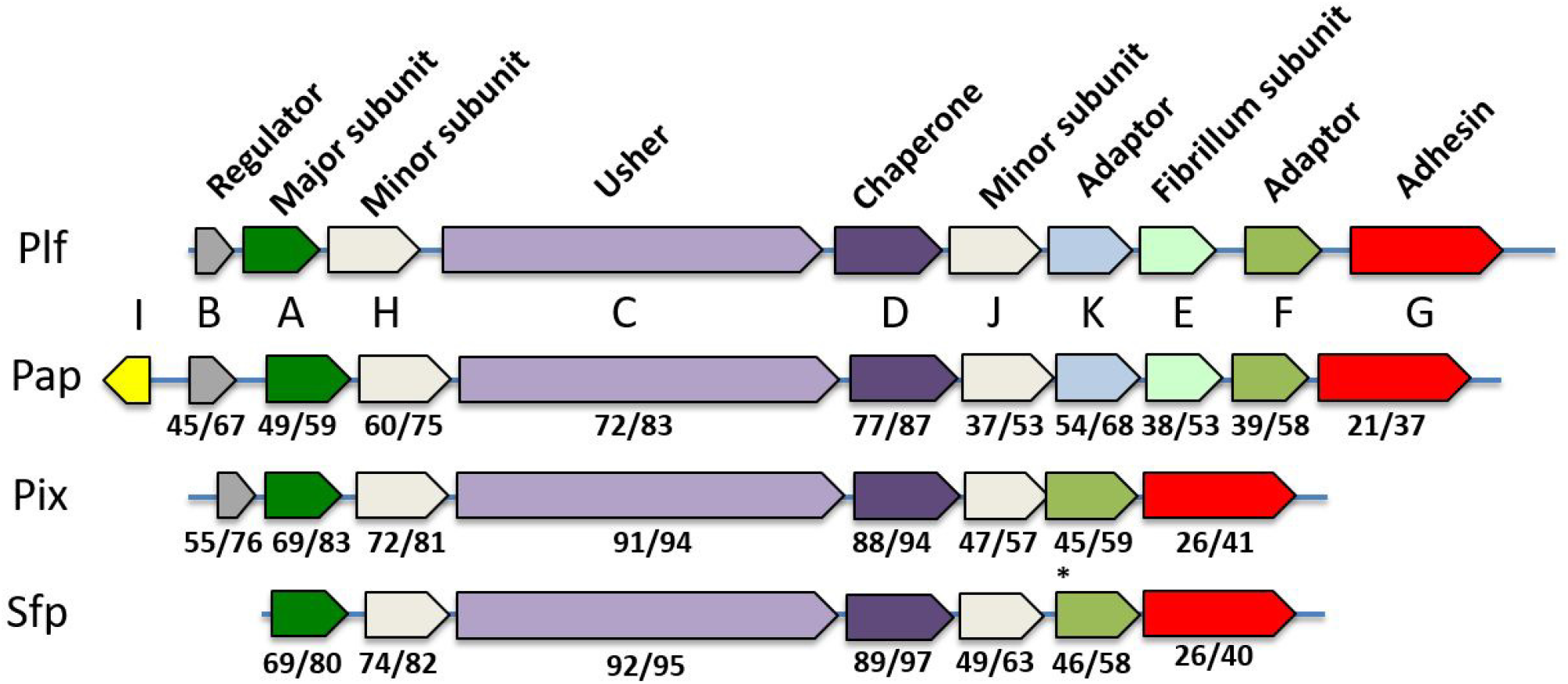
Genetic organization of the strain QT598 PL fimbrial (*plf*) gene cluster and comparison to related fimbrial gene clusters. The genes are labelled based on the *pap*- gene cluster convention. The numbers below each gene are the percentage amino acid identity/similarity obtained using BLASTP (https://blast.ncbi.nlm.nih.gov/). DNA sequences used were *plf* from strain QT598 (Genbank KP119165.1, bases 5500-14700); *pap* from UPEC 536 (Hochhut *et al*., 2006) (Genbank CP000247.1, region ECP_4533 to ECP_4543); *pix* from UPEC X2194 (Lugering *et al*., 2003) (Genbank AJ307043); *sfp* from pSFO157 (Brunder *et al*., 2001) (AF401292.1, bases 10000-17000). *SfpF was converted to a longer 186 amino acid open reading frame based on the Genbank submission. Predicted functions of gene products are indicated above. Colors denote paralogs from each fimbrial gene cluster. The *sfp* and *pix* clusters lack genes encoding PapK and PapE paralogs, that encode an adaptor controlling fibrillum length (Jacob-Dubuisson *et al*., 1993) and minor fibrillum subunits (Kuehn *et al*., 1992) respectively in P fimbriae.

### PL fimbrial gene clusters contain different types of G adhesins

To best identify potential fimbrial genes that are very closely related to the Plf system of strain QT598, alignment searches of the NCBI database were done against the predicted adhesin encoding gene product, PlfG, using a cut-off of >90% amino acid identity. The search revealed 105 samples (104 *E. coli* and one *E. albertii* strain) containing a *pflG* allele with high identity to PflG*_QT598_*. Interestingly, among the sample sources, a majority were isolated from avian species as well as clinical isolates from urine or extraintestinal infections in humans (Table S1). However, some samples were also from a variety of livestock, healthy human fecal samples, exotic zoo animal fecal samples and environmental sources. Blast analyses against PlfG*_QT598_* also identified a series of proteins demonstrating from 44% to 77% identity to PlfG that were all associated with fimbrial gene clusters belonging to the Plf family, since these fimbrial gene clusters shared the same genetic organization as the *plf* gene cluster and had highly conserved identity (>94%) with the PlfD_QT598_ gene products (data not shown). Based on sequences in the database and identification of entries containing enough sequence data to span the length of the fimbrial gene clusters, a phylogenetic analysis of distinct protein entries for different PlfG adhesins was determined. In all, 21 protein entries sharing identity with PlfG were identified (Fig. 4). Analysis determined 5 distinct clades of the PlfG adhesins, including a group (class V) specific to some *Cronobacter* spp. The number of individual entries from the sequence database indicated that PlfG class I and PlfG class II families were predominant, whereas PlfG classes III to V were represented by only a few individual strains in the sequence database (Fig. 4). All of the *plf* gene clusters identified from *E. coli* strains regardless of G adhesin class were inserted adjacent to a site-specific integrase and REP FIB region (data not shown), suggesting that these fimbrial systems are likely to be plasmid encoded. Taken together, these results suggest that *plf* gene clusters are present in numerous *E. coli* strains and that the G adhesins of these fimbriae have diversified into distinct alleles.

**Fig. 4.**
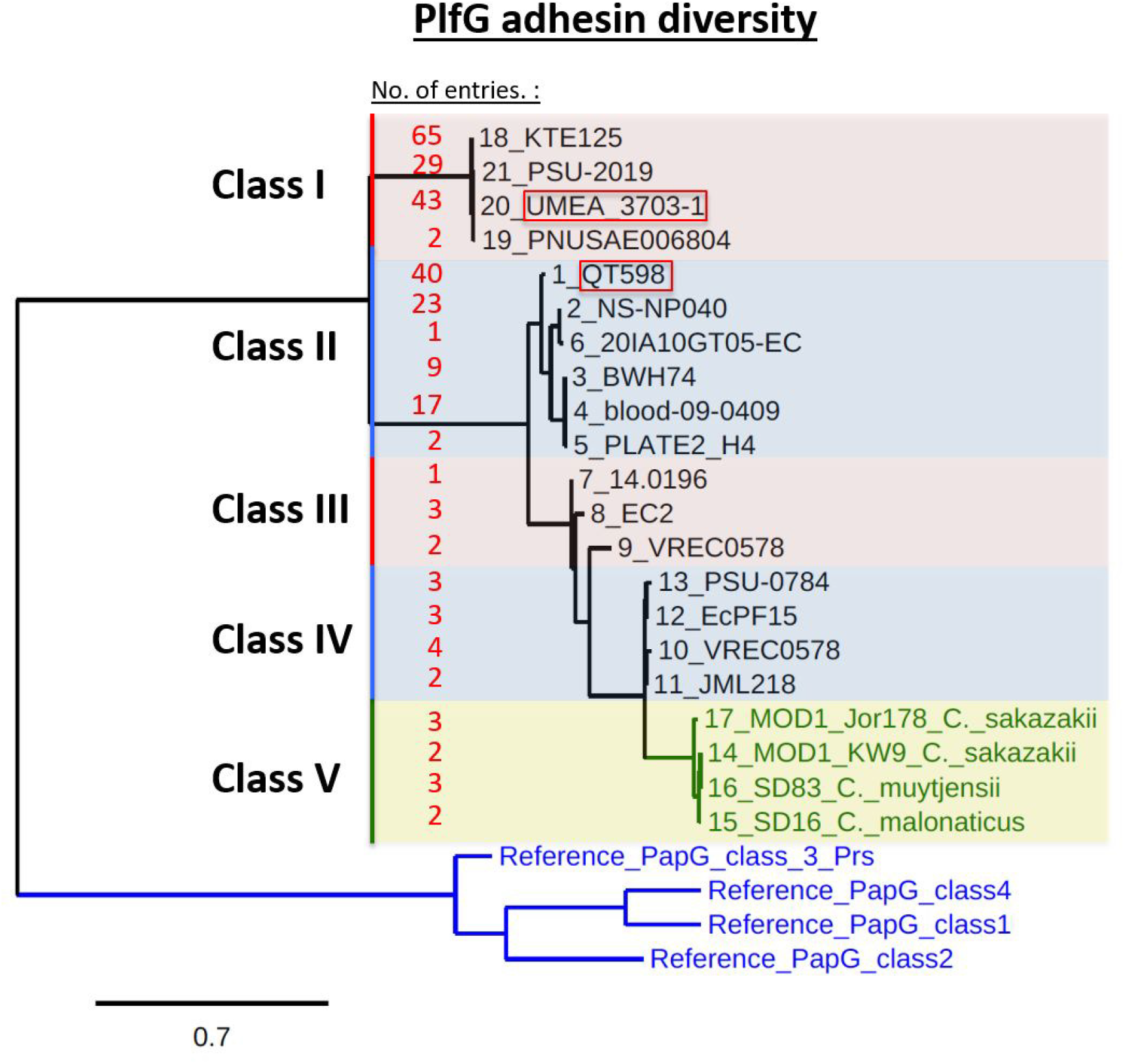
Phylogenetic analysis of different PlfG adhesin proteins. Predicted PlfG proteins from individual isolates were obtained from the sequence database at NCBI (https://www.ncbi.nlm.nih.gov) and one individual isolate sequence was selected based on sequence diversity and association with a complete *plf* fimbrial gene cluster. The total number of protein accessions for each group (No. of entries), at time of submission, are listed on the left in red. Multiple alignment (Muscle), and phylogeny (PhyML) were generated using Phylogeny.fr (www.phylogeny.fr). Analysis determined 5 distinct clades of PlfG adhesins. Including a group (class V) specific to some *Citrobacter* spp. (indicated in green). The PapG reference adhesins from P fimbriae clustered together as a distinct group from all of the PlfG adhesin proteins. The *plf* gene clusters from two strains: UMEA-3703-1 (PlfG class I) and QT598 (PlfG class II), both circled in red, were cloned for further analysis. The total number of protein accessions for each group, at time of submission, are listed on the right. Twenty-one different non-redundant entries were used: 1-WP_059331527.1; 2- WP_137488293.1; 3-WP_097732425.1; 4- WP_033555940.1; 5-WP_201475228.1; 6-EGW8442016.1; 7-MBB8123006.1; 8-WP_029305610.1; 9-WP_112039355.1; 10- WP_096965282.1 11-WP_137504062.1; 12- WP_176323703.1; 13-EFO1491433.1; 14- WP_133116004.1 15-WP_158696804.1; 16- WP_158685756.1; 17-WP_105536056.1; 18 WP_001523394.1; 19-EFB9349400.1; 20-WP_016233112.1; 21- WP_033549358.1. Alignment also included reference entries for the 4 established PapG alleles: PapG-I (strain J96)-CAA43570.1; PapG-II (strain IA2) - AAA24293.1; PapG-III (strain J96)-P42188; class PapG class IV (AAK08949.1) strain BF31.

**Table 1.**
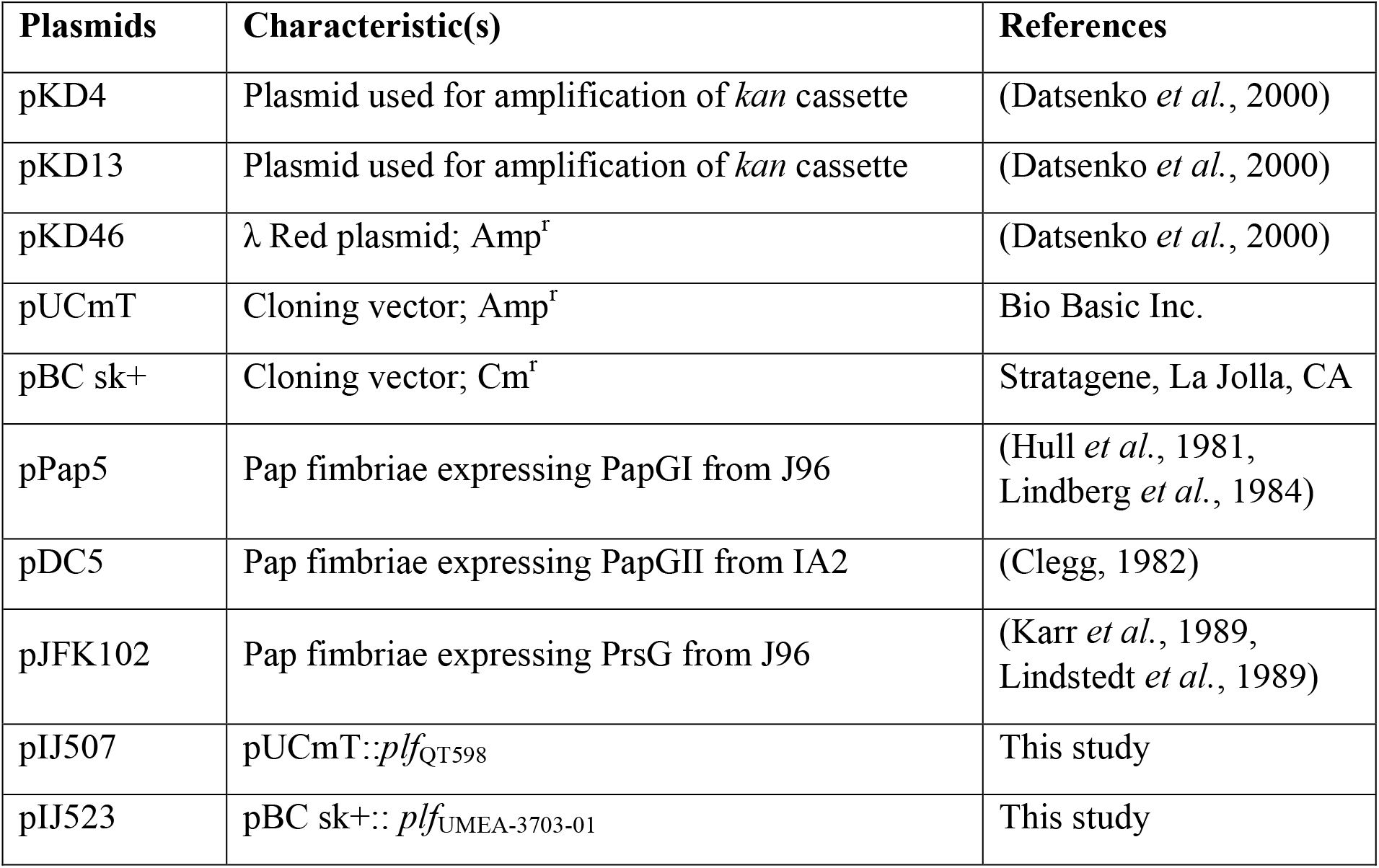
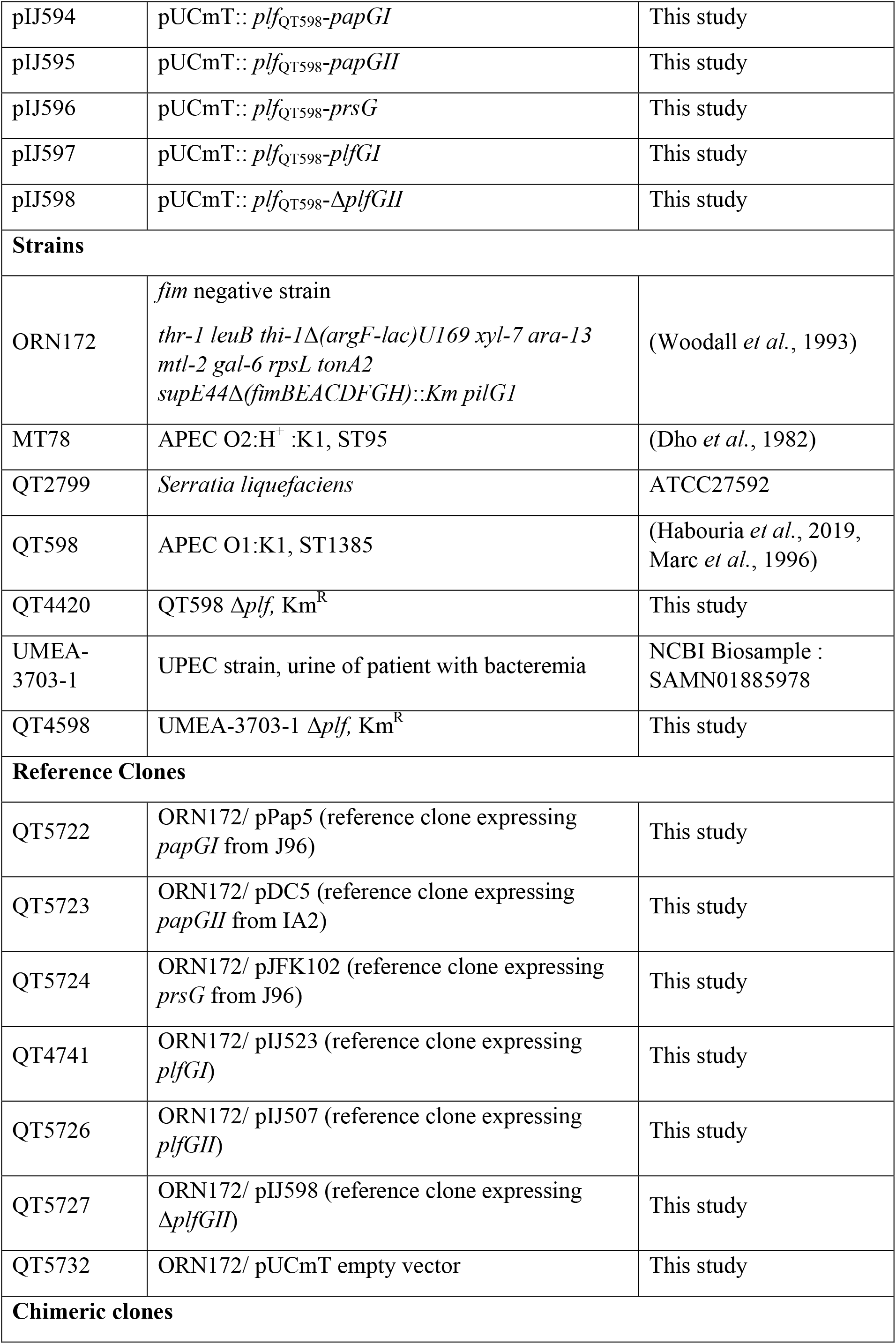
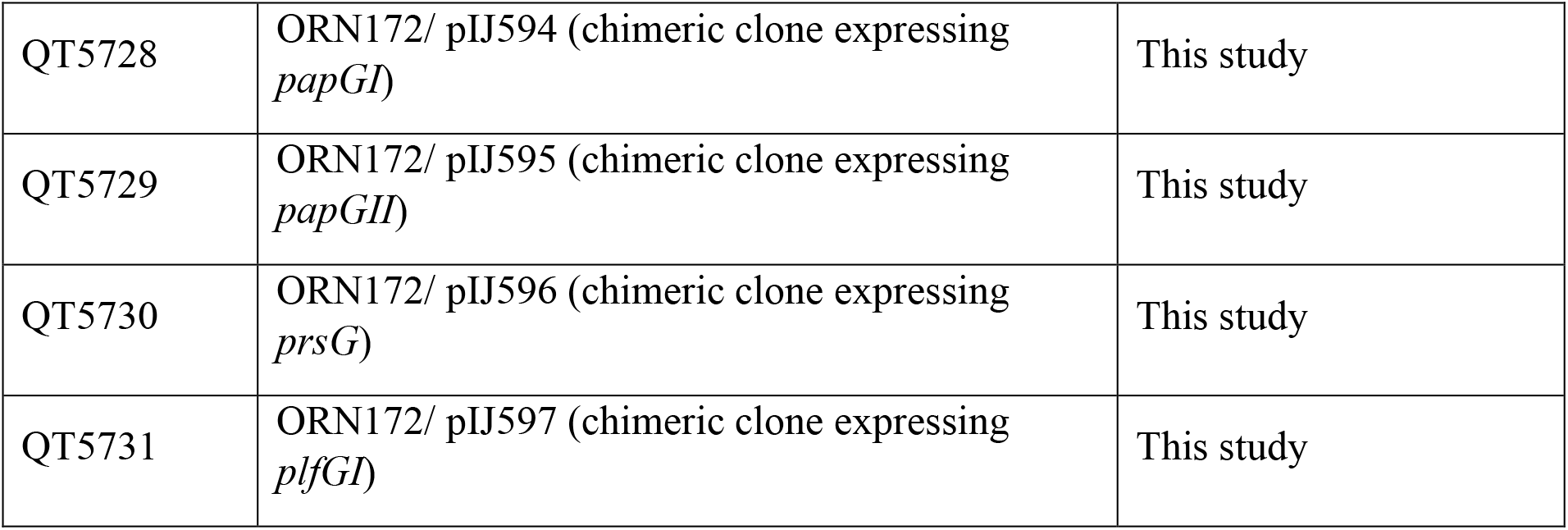
Plasmids and Strains used in this study.

Phylogenetic analysis based on the comparison of the PlfG adhesin sequences in the database demonstrated that two main classes of PlfG adhesins, class I and class II are predominant in sampled genomes. Specific Blast comparisons of the PlfG class II adhesin from *E. coli* strain QT598 with a representative encoding the Class I adhesin, from strain UMEA-3703-1, showed a 45% identity / 65% similarity. This sequence divergence is similar to the difference between P fimbriae class I and class II G adhesins (46% identity / 64% similarity). As such, and since these two PlfG classes are the most common in the database, we cloned both of them for further investigation.

### The *plf* class I and II gene clusters encode fimbriae with distinct hemagglutin activity

To demonstrate that the *plf* encoding clones produced fimbriae, the plasmids encoding *plf* genes were transformed into the afimbriated *E. coli* K-12 strain ORN172. Transmission electron microscopy (TEM) demontrated that both *plf*_QT598_ and *plf*_UMEA-3703-1_ containing plasmids produced peritrichous fimbrial filaments at the surface of cells of strain ORN172 (Fig. 5). By contrast, ORN172 containing the empty vector did not produce any fimbriae, as expected.

**Fig. 5.**
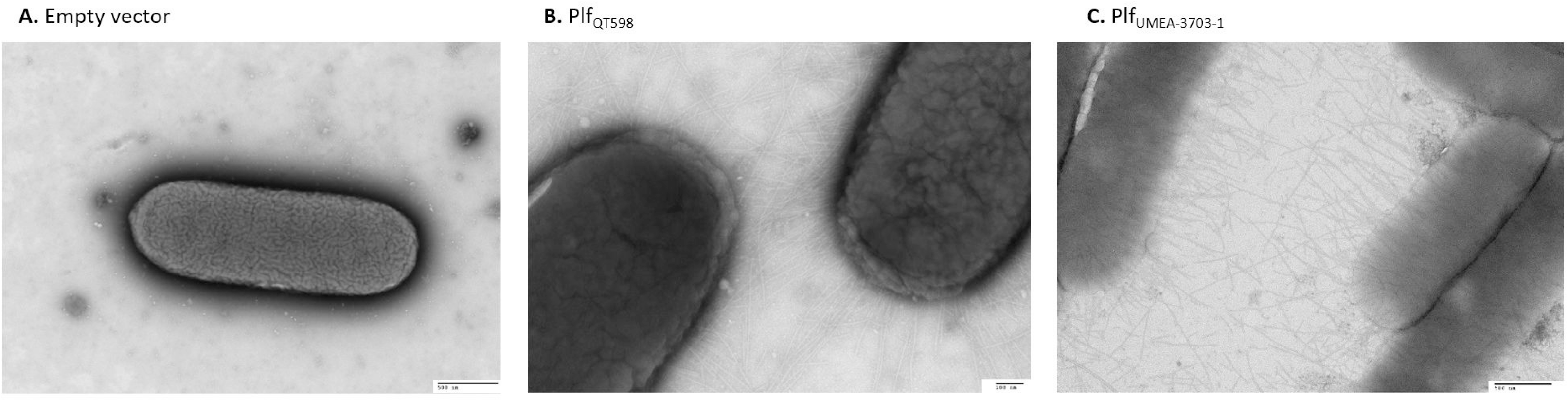
PL fimbriae visualized by transmission electron microscopy. A) ORN172 with empty vector showing no fimbriae, Bar=500 nm. B) ORN172 with plasmid pIJ507 containing *plf*_QT598_ Bar=100 nm. C) ORN172 with plasmid pIJ507 containing *plf*_UMEA3703-1_, Bar=500 nm.

P fimbrial adhesins are known to be mannose-resistant hemagglutinins and they demonstrate lectin activity specific to Gal(α1-4)Gal-containing glycolipids present on the surface of erythrocytes and other host cells. To compare the hemagglutination activity of P-fimbriae reference clones with clones producing PL fimbriae, we tested hemagglutination activity of fimbriae expressing clones in the non-fimbriated *E. coli* strain ORN172 for a variety of erythrocytes from different species (Fig. 6). The reference clone encoding P fimbriae with the PapG class I adhesin from *E. coli* J96 (pPap5) demonstrated strong hemagglutinin activities with human, pig, dog, and rabbit erythrocytes. The reference clone encoding P fimbriae containing the Pap class II adhesin from *E. coli* IA2 (pDC5) strongly agglutinated pig and human erythrocytes and, to a lesser extent, sheep and chicken erythrocytes. The reference clone encoding Prs fimbriae with the PapG class III (PrsG) adhesin from uropathogenic *E. coli* J96 (pJFK102) agglutinated dog, pig, and sheep erythrocytes. The clone encoding PL fimbriae containing a Plf class I adhesin from *E. coli* UMEA-3703-1 agglutinated a broad range of erythrocytes from all species tested except dog blood, although HA titers were higher for human and sheep blood. Interestingly, the clone encoding the Plf class II adhesin from *E. coli* QT598 only strongly agglutinated human and turkey blood (Fig. 6). Taken together, these results indicate that, as with the P fimbrial classes of adhesins, PL fimbriae are hemagglutinins and that the PlfG class I and class II adhesins demonstrate distinct hemagglutination activities compared to the P fimbrial adhesins.

**Fig. 6:**
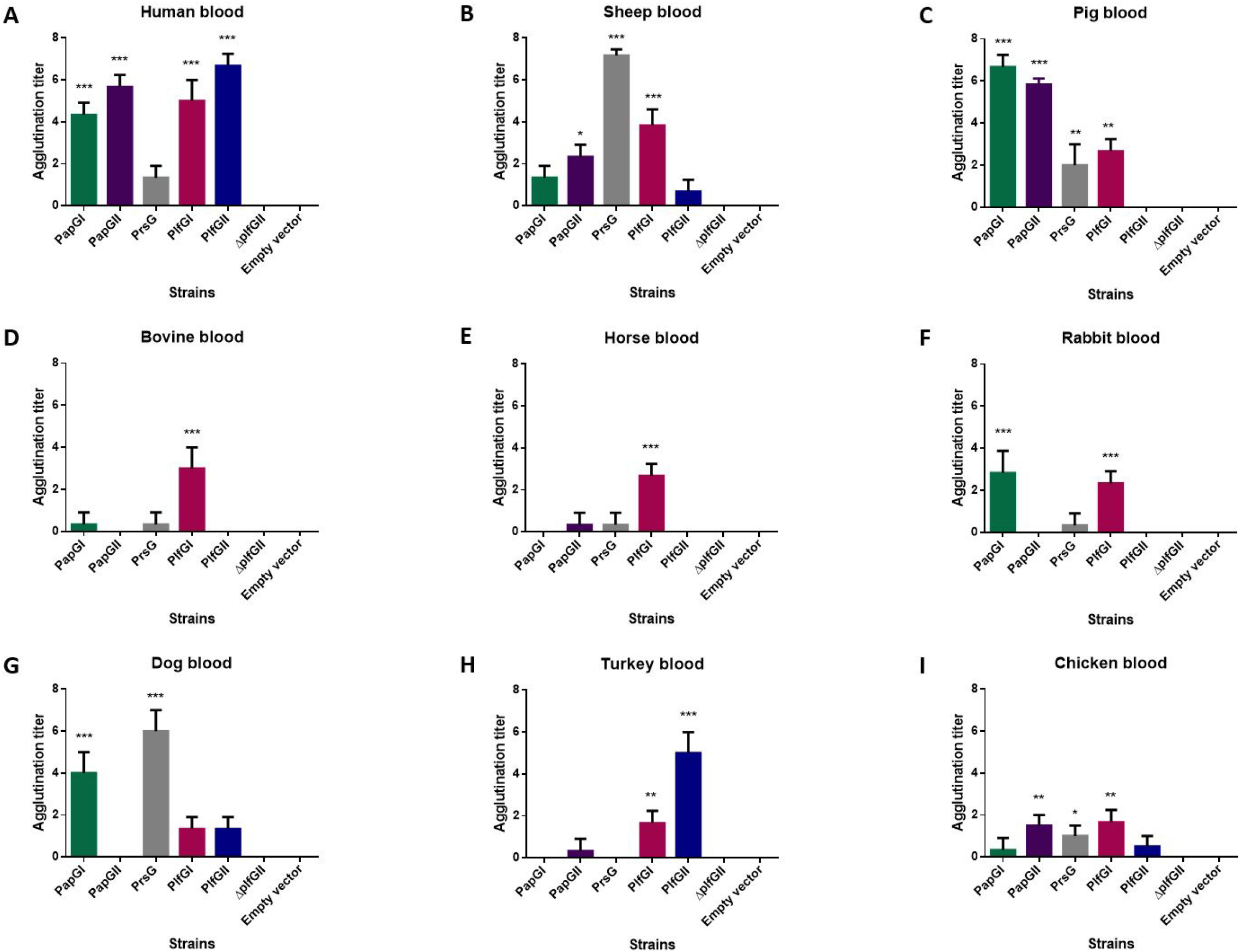
Hemagglutination activity of different clones expressing P or PL-fimbriae. Clones were *E. coli* afimbriated *fim*-negative strain ORN172 expressing P or PL- fimbriae. Cells were adjusted to an O.D._600nm_ of 60 and then diluted 2-fold in 96-well plates containing a final concentration of 3 % erythrocytes from different species. Titers are the average maximal dilution showing agglutination. Both human A and O blood gave similar titers. Reference clones showed different hemagglutination activity. However, Δ*plfG* clone as well as the empty vector showed no hemagglutination activity of any of the erythrocytes tested. (*p<0.05, **p<0.01, ***p<0.001 vs empty vector by one-way ANOVA). Plasmids used were pPap5 (*papGI*), pDC5 (*papGII*), pJFK102 (*prsG*), pIJ523 (*plf_UMEA-3703-1_*- *plf class I*), pIJ507 (*plf_QT598_*- *plf class II*), pIJ598 (*plf*_QT598_Δ*plfG*).

### Different Plf and Pap G adhesin alleles can be expressed by PL fimbriae

Since the PlfG class I and II adhesin sequences from *E. coli* strains QT598 and UMEA-3703-1 respectively are quite distinct from each other and as the *plf* gene clusters share close genetic organization with *pap* gene clusters, we generated chimeric fimbrial gene clusters encoding different G adhesin alleles. These chimeric clones were based on the *plf*_QT598_ gene cluster by generating a clone lacking the *plfG* gene (pIJ598) and then cloning the *plfG*_UMEA-3703-1_ or *papG* alleles from each of the three PapG adhesin classes (Supplemental Fig. S2. A and B). Electron microscopy demonstrated that each of the five chimeric clones introduced to non-fimbriated *E. coli* ORN172, produced fimbriae on the surface of the cells (Fig 7. A). The PlfA subunit protein was also detected from cell surface extracts as shown in immunoblots, although the level of protein present was decreased when the *papG* recombinant alleles were expressed compared to the *plfG*_QT598_ and *plfG*_UMEA-3703-1_ expressing clones (Fig. 7. B).

**Fig. 7.**
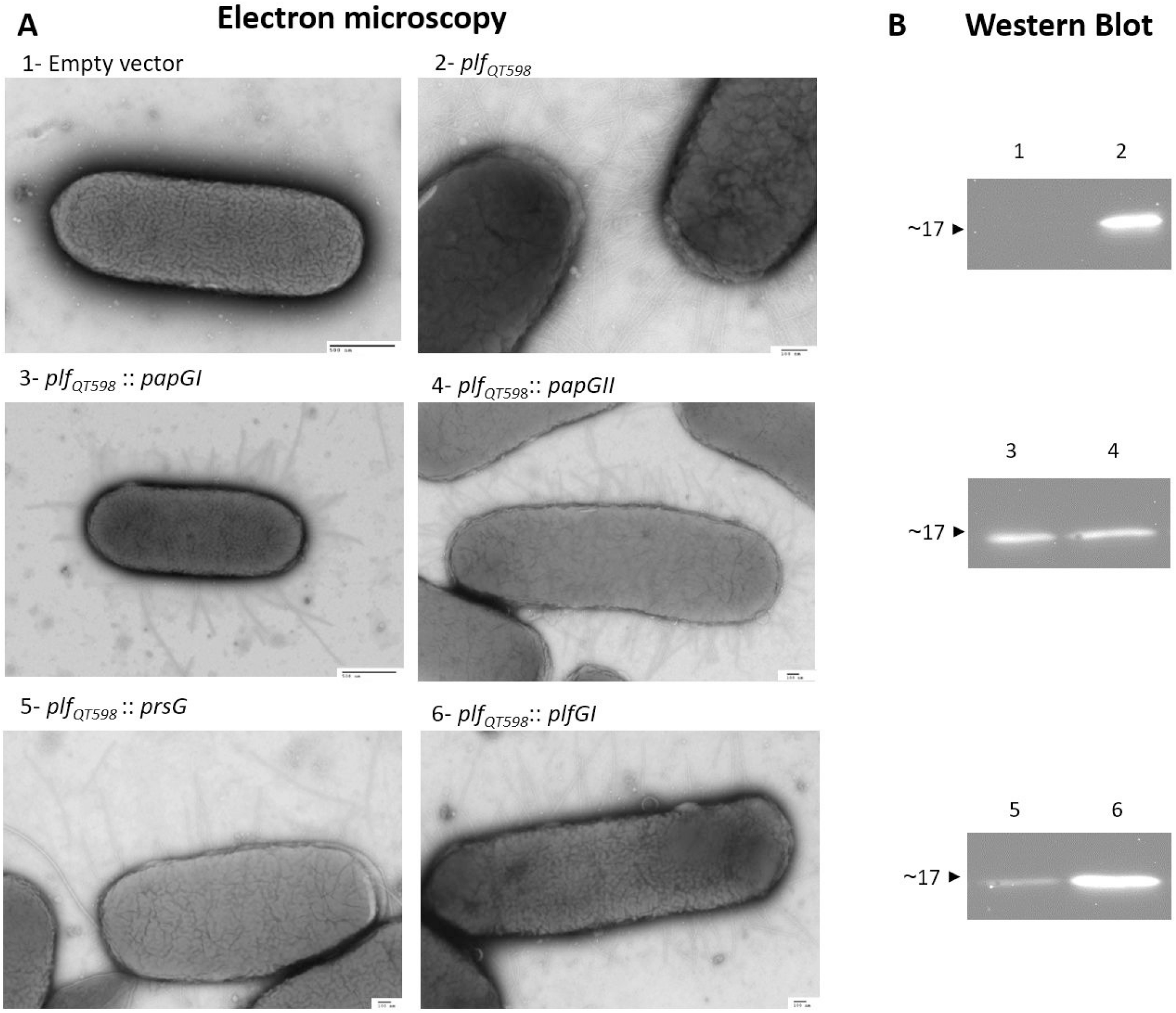
Cloning and expression of PL and Pap chimeric fimbriae. A. Chimeric clones expressing different classes of PapG/PlfG were visualized by electron microscopy. The Plf class II clone was used as a template to generate chimeric clones**. B.** Western Blot analysis of heated surface protein extracts from *plf* and chimeric clones. Antibodies used were polyclonal rabbit antibodies raised against a peptide corresponding to the fimbrial major subunit protein PlfA.

### PL fimbriae mediate adherence to human epithelial cells

The adherence of bacteria to host epithelial cells such as bladder and kidney cells is an important step in colonization of the urinary tract. To investigate whether PL fimbriae can mediate adherence to host cells, we used clones expressing PlfG class I, PlfG class II, PapG class I, II, or III fimbrial adhesins.

All the clones encoding P or PL fimbrial adhesins (reference and chimeric clones) demonstrated increased adherence to bladder 5637 and kidney HEK-293 epithelial cell lines compared to the strain containing the empty vector (Fig. 8). The chimeric clones containing the *plf* gene cluster with hybrid *pap* or *plfG* adhesin encoding genes also promoted adherence to epithelial cells. However, the clone containing a *plf* gene cluster, lacking the *plfG* or *papG* gene showed no appreciable adherence compared to the empty vector-containing clone (Fig. 8). These results demonstrate that PL fimbriae producing distinct types of G adhesins can mediate adherence to urinary tract epithelial cells, and suggest that these fimbriae may potentially play a role during host colonization.

**Fig. 8.**
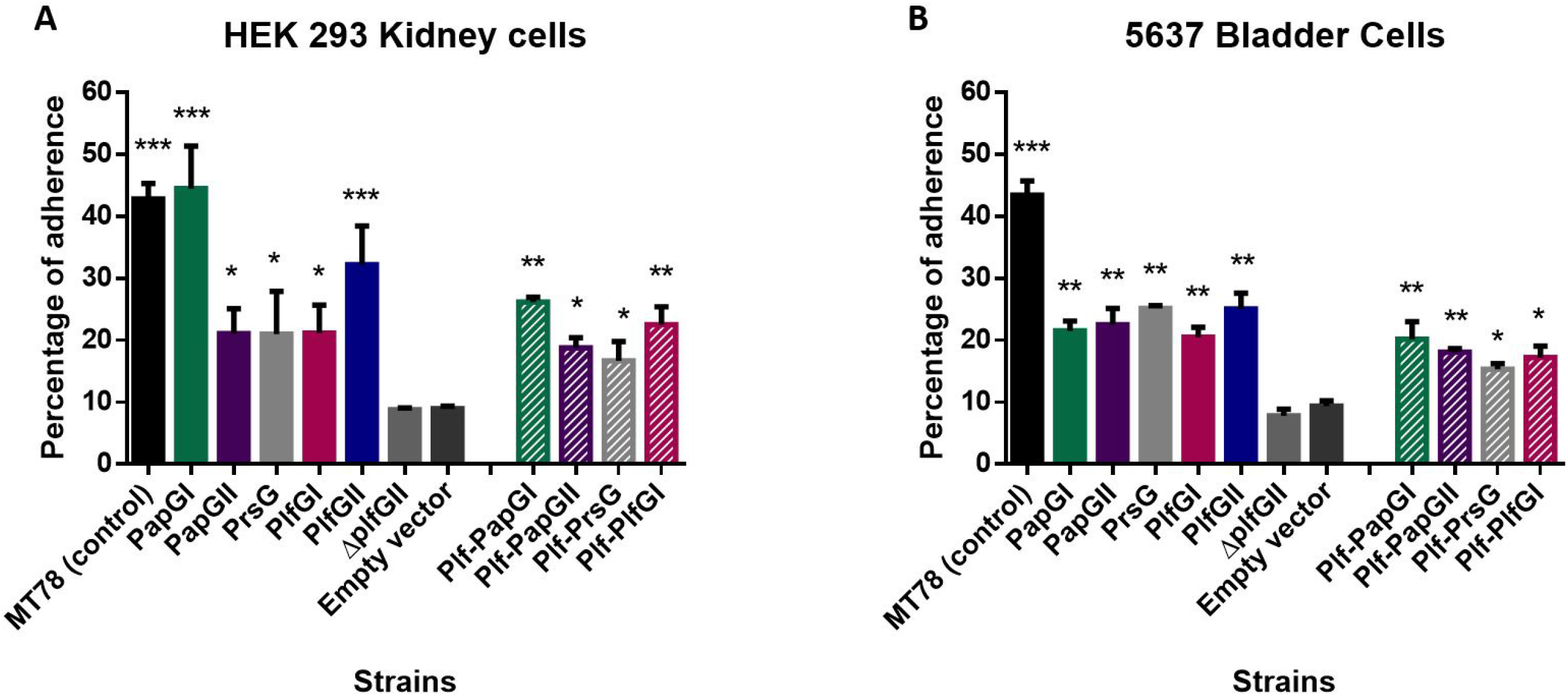
Reference and chimeric clones promote adherence to human kidney (HEK-293) and bladder (5637) epithelial cells. Cell monolayers were infected with *E. coli fim-*negative ORN172 expressing P and PL fimbrial proteins at a multiplicity of infection (MOI) of 10 and incubated at 37°C at 5% CO_2_ for 2 hours. Adherent bacteria were enumerated by plating on LB agar. Empty vector was used as a negative-control and APEC MT78 as a positive control for adherence to cell lines. All the clones encoding P or PL fimbrial adhesins (reference and chimeric clones) demonstrated increased adherence to human bladder 5637 and kidney HEK-293 epithelial cell lines compared to the strain containing the empty vector. The Δ*plfG* clone also did not adhere to human epithelial cells. Data are the averages of three independent experiments. Error bars represent standard errors of the means (*p<0.05, **p<0.01, ***p<0.001 vs empty vector by one-way ANOVA).

### PL fimbriae promote biofilm production

Since fimbriae can contribute to biofilm production, we tested for biofilm formation in polystyrene microtiter plates at different temperatures (25°C, 37°C, and 42°C) (Fig. 9). The clone expressing PL fimbriae with the PlfG class II adhesin showed a high level of biofilm production at all tested temperatures, even above that of a positive control biofilm forming *Serratia liquefaciens* (*S. liquefaciens*) reference strain. The clone expressing PL fimbriae with the PlfG class I adhesin also produced biofilm at 25°C and 37°C at moderate levels compared to the clone producing the PlfG class II adhesin. However, biofilm production was very low at 42°C. The Pap class I and III producing reference clones were also able to form significantly more biofilm at 25°C, 37°C, and 42°C than the negative control. The Pap class II expressing reference clone produced biofilm at higher temperatures (37°C and 42°C), but biofilm levels were reduced at 25°C.

**Fig. 9.**
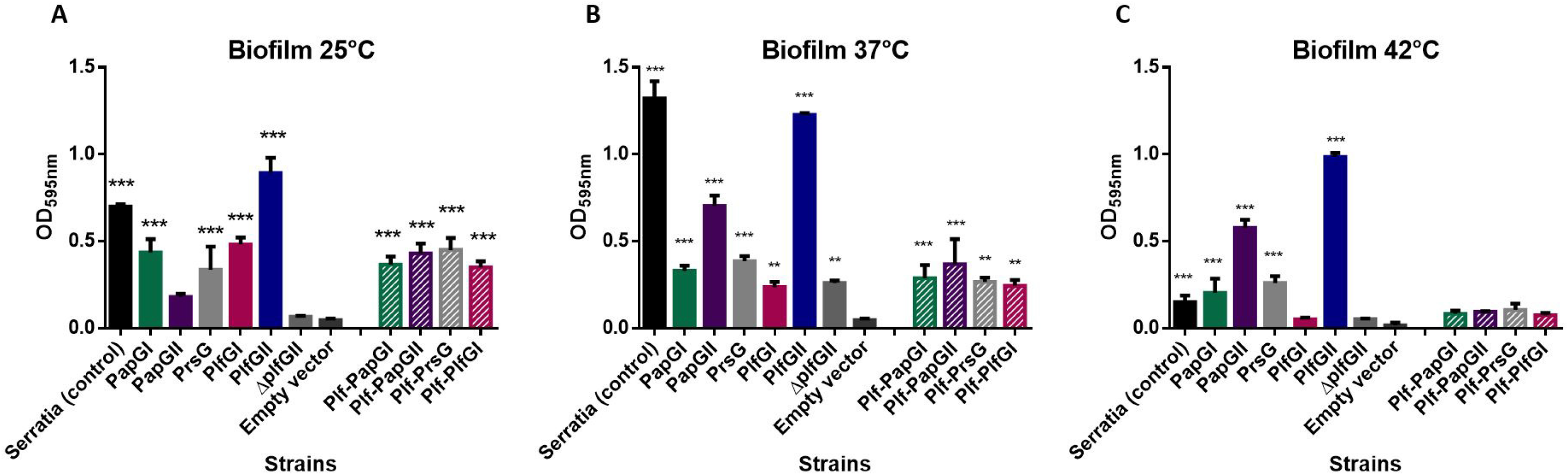
Biofilm production by clones expressing Plf_QT598_ and Plf_UMEA-3703-1_ and reference and chimeric clones at different temperatures. Clones of *E. coli* strain ORN172 expressing P and P-like fimbriae proteins were grown at different temperatures (25, 37 and 42°C) in polystyrene plate wells for 48 hours and then stained with crystal violet. Remaining crystal violet after washing with acetone was measured as absorbance at 595nm. Data are the means of three independent experiments and error bars represent standard errors of the means (*p<0.05, **p<0.01, ***p<0.001 compared to empty vector using one-way ANOVA). Empty vector was used as a negative-control and *S. liquefaciens* strain was used as a positive control for biofilm production.

The chimeric clones that expressed different Pap or Plf adhesins fused to the *plf*_QT598_ gene cluster were all able to produce appreciable levels of biofilm at both 25°C and 37°C, although biofilm was much reduced at 42°C. The clone expressing the *plf*_QT598_ gene cluster lacking a *papG* or *plfG* adhesin encoding gene (Δ*plfG* clone) as well as the empty vector were not able to produce biofilm at all the tested temperatures (Fig. 9). Taken together these results indicate that PL and P fimbriae expressing different types of G adhesins can mediate biofilm production in *E. coli* K-12 and that the PlfG class II adhesin in particular can contribute to strong biofilm formation.

### PL fimbrial genes are upregulated in the bladders of infected mice

To investigate the potential role of the PL fimbriae in virulence in the UTI model, 6- week-old female CBA/J mice were infected with wild type strains QT598 and UMEA-3703-1 or with mutant Δ*plf* strains, lacking the genes encoding PL fimbriae. In the mouse infection model, loss of PL fimbriae did not have a significant affect on colonization of the bladder or kidneys by strain QT598 (Fig 10.A). Strain UMEA-3703-1 and its Δ*plf* mutant also showed no significant differences in colonization. However, UMEA-3703-1 was only able to colonize at lower levels (10^2^ to 10^3^ cfu/g) compared to strain QT598 (10^5^ to 10^6^ cfu/g) (Supplemental Fig. S3). Interestingly, the expression level of *plf*_QT598_ was upregulated by more than 40-fold in the bladder of infected mice when compared to expression following growth in vitro in LB medium (Fig 10.B). This suggests that the expression of this fimbriae is favored by environmental cues during infection in the urinary tract.

**Fig. 10.**
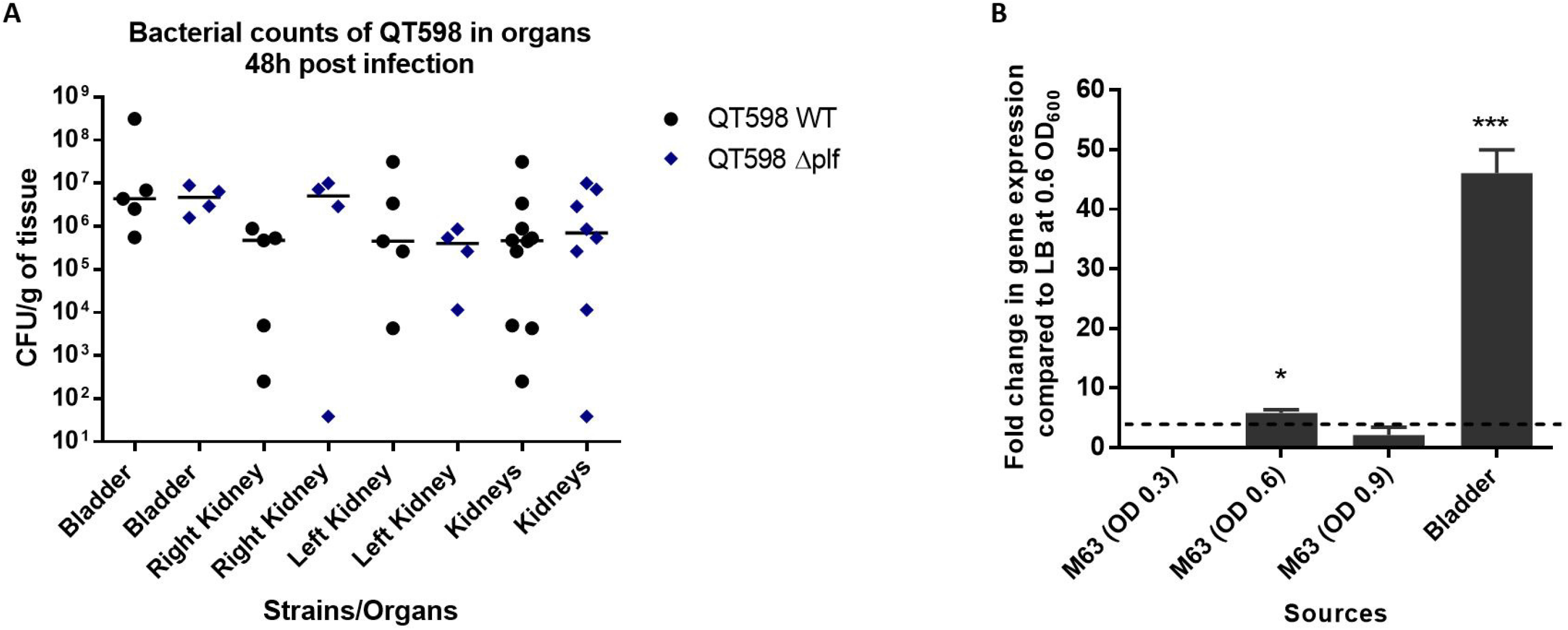
Loss of PL fimbriae in the murine model of ascending UTI does not reduce colonization. CBA/J mice were infected with either the WT strain QT598 or an isogenic Δ*plf* mutant (Δ*plf*). Mice were euthanized after 48h, and bladder and kidneys were harvested for colony counts. **A.** Infections were performed to compare wild-type strain QT598 to its mutant. There were no significant differences in colony counts in either bladders or kidneys (Data are means +/- standard errors of the means. * p < 0.05, ** p < 0.01, Mann-Whitney Test). **B.** RT-PCR analysis of *plf* expression by strain QT598. QT598 was grown in LB medium to OD_600_ of 0.6 and used as a standard to compare it with growth in M63 minimal medium (with glycerol as carbon) at different growth phases (OD_600_ of 0.3, 0.6. and 0.9). RNAs were also extracted from infected bladders. Transcription of *plf* was significantly upregulated in the mouse bladder. (*p< 0.05, ***p<0.001, error bars indicate standard deviations, Student t-test). The dashed line corresponds to the cutoff for a significant difference in expression.

## Discussion

A novel plasmid-encoded fimbrial gene cluster was identified on the large colicin V plasmid of avian pathogenic *E. coli* strain QT598 (Serotype O1:K1, Sequence type ST1385). Strains from this and related sequence types such as ST91 are commonly associated with extra-intestinal infections in poultry and urinary tract infections in humans (Habouria *et al*., 2019) and (http://enterobase.warwick.ac.uk/). The *plf* genes were shown to be adjacent to the repFIB and *intM* genes on plasmid pEC598. This is a common integration site for a diversity of genes on F and related plasmids, and collectively this region has been named the « cargo gene » region (Koraimann, 2018). The cargo gene region has been found to encode a diversity of accessory genes, insertion sequences, and integrons known to encode genes for resistance to antimicrobials and metals, microcins, and virulence genes (Koraimann, 2018, Lanza *et al*., 2014). It is therefore likely that the *plf* fimbrial gene cluster along with other genes was inherited by certain *E. coli* strains through a recombination/integration event and that it has since disseminated or been transferred into a diversity of *E. coli* isolates associated with different host or environmental sources (highlighted in Table S1). As with the P fimbriae, PL fimbriae have also diversified considerably, and there has been notable divergence in the sequence of the PlfG adhesin encoding sequences into 5 distinct PlfG adhesin classes (Fig. 4).

Such changes may have occurred to promote adherence and colonization to a variety of surfaces or host cell receptors in different niches or environments.

The PL fimbriae are new members of the π fimbrial family, which contains P-fimbria- like operons present in some Betaproteobacteria and Gammaproteobacteria (Nuccio *et al*., 2007). More specifically, based on comparison of the fimbrial usher proteins, the PL fimbriae are part of a subgroup which includes true P fimbriae, as well as closely related Sfp and Pix fimbriae (Nuccio *et al*., 2007) (Fig. 2), all of which have been shown to mediate mannose-resistant hemagglutination (MRHA) of erythrocytes from humans in addition to some distinct MRHA profiles for erythrocytes from other species. Pix fimbriae, which have been identified in some uropathogenic *E. coli* strains, were shown to agglutinate human erythrocytes, but not sheep or goat erythrocytes and do not recognize the Gal-Gal sugars recognized by P fimbriae (Lugering *et al*., 2003). Sfp fimbriae also mediate MRHA of human erythrocytes, which was dependent on the *sfpG* gene (Brunder *et al*., 2001). However, to our knowledge, no tests for MRHA for erythrocytes from other species have been reported. Interestingly, the G adhesin proteins from Pix and Sfp fimbriae share amino acid homology between them (63% identity/81% similarity), suggesting these G adhesin proteins are more closely related to each other than to PlfG or PapG adhesins, which share no more than 25% amino acid identity. Herein, we demonstrated that PL fimbriae producing the class I adhesin mediated MRHA for erythrocytes from different species including equine, ovine, bovine, rabbit and human erythrocytes, whereas PL fimbriae producing the class II adhesin mediated MRHA only to human and turkey erythrocytes (Fig. 6). Taken together, this subgroup of π fimbriae (true P fimbriae, Sfp, Pix, and PL fimbriae) have developed important differences in adhesin protein sequences that have expanded the capacity to adhere to a variety of receptors on erythrocytes and host cells from different species. It will be of interest to more specifically determine the lectin receptor specificity of this family of fimbriae.

The genetic organization of the *plf* gene cluster includes 9 predicted fimbrial subunit genes, which is the number of predicted structural genes encoding P fimbriae (Fig. 3). By contrast, both the Sfp and Pix fimbrial gene clusters comprise 7 structural genes, and lack the genes corresponding to the *papK* and *papE* genes predicted to encode an adaptor and a minor fimbrial subunit (Fig. 3). From this standpoint, overall, PL fimbriae are most similar to true P fimbriae.

To further demonstrate potential complementarity between P and PL fimbriae, we also generated hybrid fimbrial gene clusters, wherein the *plfG*_QT598_ gene was replaced by PapG adhesin encoding genes belonging to class I, class II or class III adhesins. Each of these clones were able to produce functional fimbrial structures that also increased adherence to human urinary tract epithelial cells. This also further indicates that the PL fimbriae, despite having adhesins that are quite distinct in amino acid sequence from P fimbriae, also produce mannose-resistant hemagglutinins that can mediate adherence to human bladder and kidney cells, and that the bioassembly of these fimbriae are compatible with P fimbrial G adhesins. It is interesting, however, that the production of the hybrid fimbriae from bacterial cells was substantially reduced compared to the PL fimbrial clones, suggesting that efficiency of biogenesis of the hybrid fimbriae is reduced.

As with the *plf* gene cluster, the location of the *sfp* genes is also on IncF plasmids, in close proximity to the repFIB region on the pSFO157 plasmid (Brunder *et al*., 2001). However, it is flanked on both sides by insertion sequences that are distinct from the region adjacent to *plf* genes on pEC598. The Sfp fimbriae were initially found to not be expressed by EHEC strains under normal laboratory conditions, and properties of these fimbriae were first determined using cloned fimbrial genes in *E. coli* K-12 (Brunder *et al*., 2001). The *sfp* genes encoding a fimbrial system with mannose-resistant hemagglutinin activity have been identified on a subgroup of sorbitol-fermenting EHEC/STEC strains and some EHEC O165:H25/NM strains from humans and cattle, but are absent from most other types of *E. coli* (Bielaszewska *et al*., 2009, Brunder *et al*., 2001). This suggests that the *sfp* genes were likely acquired independently by horizontal transfer to both a non-motile sorbitol O157 strain and independently to an O165:H25/NM strain and have since remained in these branches of EHEC (Bielaszewska *et al*., 2009). This is clearly in contrast to the *plf* gene cluster, which is present in a diversity of *E. coli* strains from multiple sources, and has likely been transferred either through multiple conjugation and/or recombination events and has also diversified, since distinct G adhesin classes have emerged among strains.

DNA sequence comparisons of gene clusters that are highly similar to the *plf* fimbrial system of *E. coli* QT598 from nucleotide databases provided a means to identify subgroups of PL fimbriae encoding 5 distinct classes of PlfG adhesins (Fig. 4). Since the PlfG class I and class II encoding alleles were predominant among isolates that notably included strains associated with human extraintestinal infections as well as infections from poultry, we focused our attention on functional characterization of one of each of the PL fimbriae belonging to these classes. It was also interesting to identify some variant alleles of the PlfG adhesin in other *E. coli* strains as well as a subgroup that was identified in some strains of *Cronobacter sakazakii* and other *Cronobacter* spp. (Fig 4). Although *Cronobacter* strains containing the *plf* fimbrial clusters were sampled from spices, *Cronobacter sakazaki* and related *Cronobacter* spp. are important foodborne pathogens that can contaminate dehydrated milk and other products and cause serious extraintestinal infections, particularly in neonates (Healy *et al*., 2010, Lee *et al*., 2019).

The capacity of PL fimbrie to form biofilms at different temperatures was also investigated, and both the class I and class II PL fimbriae promoted biofilm formation with PlfG class I producing more biofilm at 25°C and 37°C, but not 42°C. By contrast, the PlfG class II adhesin produced very high levels of biofilm at all temperatures tested. Presence of the PlfG adhesin was important for high-level biofilm production for PlfG class II, since the absence of the *plfG* adhesin gene greatly reduced biofilm formation. Notably, after growth at 37°C, the level of biofilm produced by the *ΔplfG* clone was significantly higher than the empty vector and comparable to levels of biofilm produced by Pap reference clones and the PlfG class I clone. This suggests that other factors in addition to the G adhesin may also contribute to increased biofilm formation associated with expression of *plf* or *pap* fimbrial genes.

Since both types of PL fimbriae conveyed increased adherence to human epithelial bladder and kidney cells (Fig. 8), we investigated the potential of these fimbriae to contribute to urinary tract colonization in a murine model. Deletion of the *plf* genes from either *E. coli* strain QT598 or strain UMEA-3703-1 did not have an appreciable effect on colonization of either the bladder or the kidneys. Further, despite being isolated from a human UTI, strain UMEA-3703-1 was not a strong colonizer in the mouse UTI model.

The mouse UTI model may not be as representative of a human infection when using certain bacterial strain backgrounds or when investigating specific mechanisms of virulence such as adherence and fimbrial adhesins. P fimbriae have been shown to play a role in urinary infection, particularly for pyelonephritis in cynomolgous monkeys (Roberts *et al*., 1994) and these fimbriae alone can confer an asymptomic *E. coli* urinary strain the capacity to elicit strong regulatory modulation in humans by acting as an IRF-7 agonist and reprogramming the immune response in the urinary tract (Ambite *et al*., 2019). In the case of the murine model, it has been demonstrated that P fimbriae can reduce the immune response in the kidney by decreased production of polymeric Ig receptor and reduced secretion of IgA (Rice *et al*., 2005). However, the role of P fimbriae in bacterial colonization in the UTI mouse model has been less evident. Initially, *pap* genes cloned into avirulent *E. coli* K-12 or an intestinal commensal *E. coli* were shown to increase colonization of the mouse kidney (Hagberg *et al*., 1983, O’Hanley *et al*., 1985). By contrast, deletion of *pap* from different UPEC strains did not alter colonization of the urinary tract in CBA/J mice (Mobley *et al*., 1993). Reasons why PL fimbriae as much as P fimbriae may not play a critical role in the mouse UTI model may be due to differences in lectin-receptor target specificity present on murine cells and/or the potential redundancy of adherence mechanisms due to production of multiple fimbrial adhesins in UPEC strains. Despite not playing a role in the UTI murine infection model, expression of the *plf* was upregulated more than 40-fold in the bladder and was increased by 5-fold in minimal medium compared to rich medium (Fig. 10.B). This indicates that the expression of these fimbriae can be increased by cues during infection, which may include host factors or decreased nutrient availability. It will be of interest to determine whether PL fimbriae or specific PlfG adhesins may contribute to infection in other animal models such as poultry and to further investigate PL fimbriae receptor specificity, potential role in modulation of host immune response and regulation of production of this newly identified group of fimbriae.

## Materials and Methods

### Bacterial strains, plasmids, and growth conditions

Bacterial strains and plasmids used in this study are listed in Table 1. ExPEC strain QT598 (a passaged derivative of strain MT156 (Marc *et al*., 1996)) is an O1:K1 sequence type ST1385 strain originally isolated from a turkey suffering from colibacillosis in France (Habouria *et al*., 2019). UPEC strain CFT073 was isolated from the blood and urine of a woman suffering from urinary tract infection (Mobley *et al*., 1990), and UMEA-3703-1 (NCBI Biosample: SAMN01885978) was isolated from the urine of a human with bacteremia. *E. coli* K-12 laboratory strains DH5α and ORN172 (type 1 fimbriae *fim*-negative strain) were used for cloning and protein expression. Reference clones expressing fimbriae encoding different PapG adhesin classes were used as controls including plasmids pPap5 (*papG*_J96_)-class I (Hull *et al*., 1981, Lindberg *et al*., 1984), pDC5 (*papG*_IA2_)-class II (Clegg, 1982), and pJFK102 (*prsG*_J96_)-class III (Karr *et al*., 1989, Lindstedt *et al*., 1989).

Bacteria were routinely grown at 37°C on solid or liquid Luria–Bertani LB medium (Alpha Bioscience, Baltimore, MD). When required, antibiotics were added to a final concentration of 100µg/ml of ampicilin, 30 µg/ml of chloramphenicol, or 50 µg/ml of kanamycin.

### Bioinformatics analysis

Identification and comparison of sequences in the databases was achieved by accessing data on completed genomes and BioProjects publicly available in the NCBI database (www.ncbi.nlm.nih.gov). Analyses included BLAST against both nucleotide and protein entries. Figures presenting the organization and comparison of genes and gene clusters were generated from the nucleotide accession numbers and entries using SnapGene (Version 5.2.1) (www.snapgene.com). For comparison of the protein sequences, entries were obtained either from NCBI or the Universal Protein Resource (UniProt) (www.uniprot.org) websites. Phylogenetic analyses of protein sequences were done using the platform at Phylogeny.fr (http://www.phylogeny.fr) using the default (“one click”) parameters (Dereeper *et al*., 2008). Analyses consisted of Multiple sequence alignment with MUSCLE (Edgar, 2004), alignment curation with GBlocks (Castresana, 2000), maximum-likelihood phylogeny analysis using PhyML 3.0 (Guindon *et al*., 2010), and TreeDyn for generation and editing of trees (www.treedyn.org). Specific parameters are described at the Phylogeny.fr website platform.

### Construction of plasmids

Cloning of the *plf* gene clusters and *plfG* and *papG* genes encoding the different classes of adhesins were obtained by PCR amplification using specific primers (Table 2) and Q5 High Fidelity-DNA polymerase (New England Biolabs [NEB]). The A-Tailing Kit (NEB) was then used to add additional deoxyadenosine (A) to the 3’ end of the PCR products. The insert possessing the additional A at 3’ end was ligated to the linearized vector with additional deoxythymidine (T) residues using T4 DNAligase (NEB). The *plf* gene cluster from strain QT598 (*plf*_QT598_) was amplified using primers CMD1847_F and CMD1900_R and cloned into vector pUCm-T (Bio Basic, Markham, Ontario, Canada), generating plasmid pIJ507. This plasmid encodes the full *plf* gene cluster with a PlfG class II adhesin. The *plf* gene cluster from strain UMEA-3703-1 (*plf*_UMEA_) was amplified using primers CMD2119_F and CMD2120_R and cloned into vector pBC sk+, generating plasmid pIJ523. This plasmid encodes the full *plf*_UMEA_ gene cluster with a PlfG class I adhesin. To generate chimeric gene clusters comprised of *plf*_QT598_ with different types of G adhesin encoding genes, pIJ507 was used as a template. *plfG*_QT598_ was deleted using an inverse PCR method with primers CMD2168_F and CMD2169_R which introduced PmeI sites and amplified a linear fragment lacking the *plfG*_QT598_ gene. The linearized product was then treated with DpnI endonuclease (NEB) to cleave any methylated template sequence. The linear fragment was either ligated using T4 DNA ligase (NEB) to generate pIJ598, which encodes *plfBAHCDJKEF* (*plf*_QT598_Δ*plfG*), or used as a template to generate chimeric fimbrial gene clusters containing different G adhesins using the T4 DNA ligase (NEB). PCR fragments containing G adhesin genes were obtained using primer pairs CMD2171_F and CMD2172_R (for *papG* class I from strain J96); CMD2174_F and CMD2175_R (for *papG* class II from strain CFT073); CMD2177_F and CMD2178_R (for *prsG* [*papG* class III] from strain J96); and CMD2180_F and CMD2181_R (for *plfG*_UMEA-3703-1_ plfG class I from strain UMEA-3703-1). Cloning experiments to generate recombinant plasmids or subclones were first achieved using *E. coli* strain DH5α. The plasmids were extracted using a Miniprep kit according to the manufacturer’s recommendations (Bio Basic Inc.) and then transformed into *E. coli fim*-negative strain ORN172 for phenotypic testing. Strains containing reference plasmids that contain full P fimbrial gene clusters were used as reference controls: pPap5 (encoding P fimbriae PapG class I from *E. coli* J96) (Hull *et al*., 1981, Lindberg *et al*., 1984), pDC5 (encoding P fimbriae PapG class II from strain IA2) (Clegg, 1982), and pJFK102 (encoding Prs fimbriae PrsG (PapG class III) from *E. coli* J96) (Karr *et al*., 1989, Lindstedt *et al*., 1989).

**Table 2.**
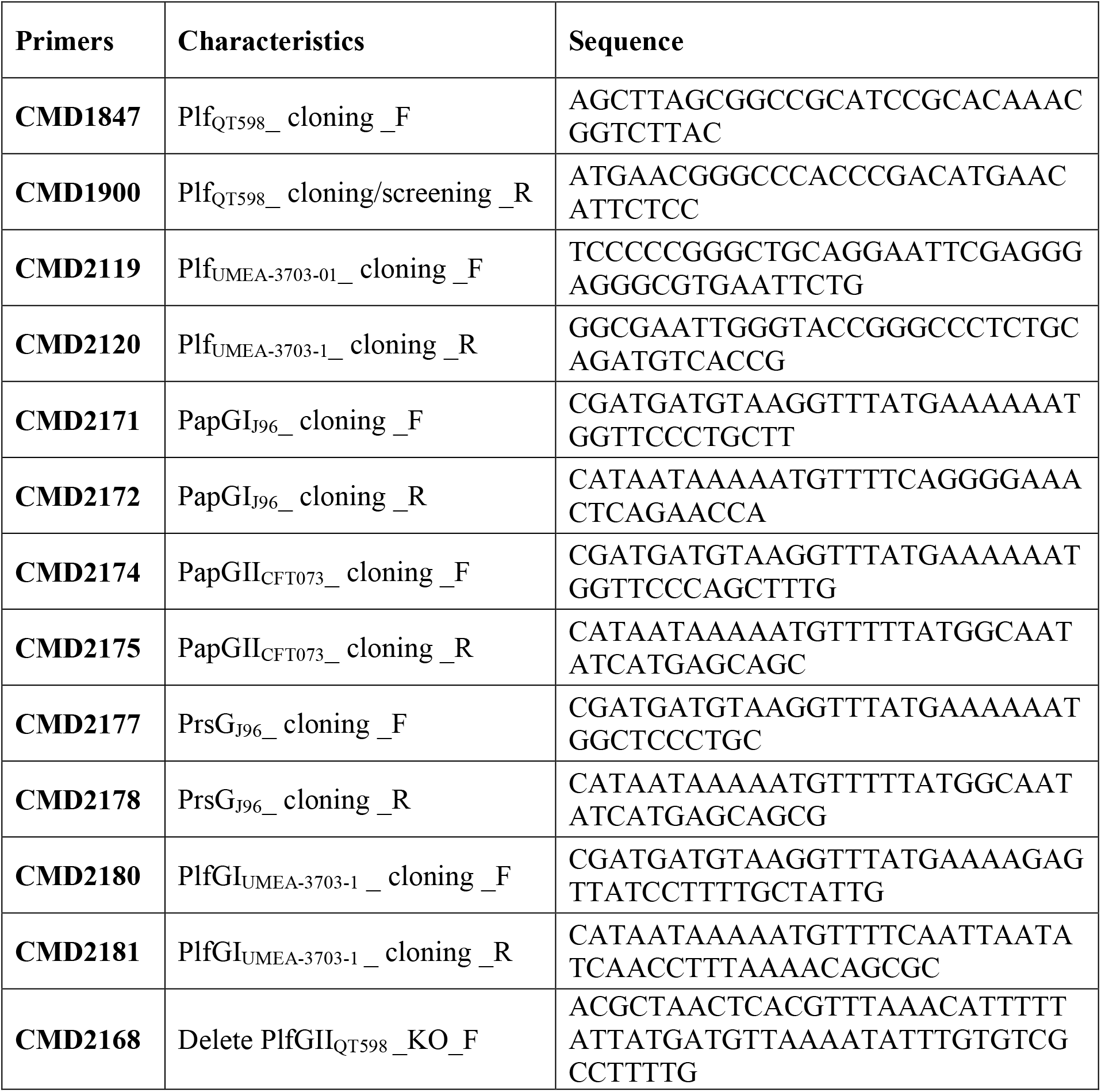
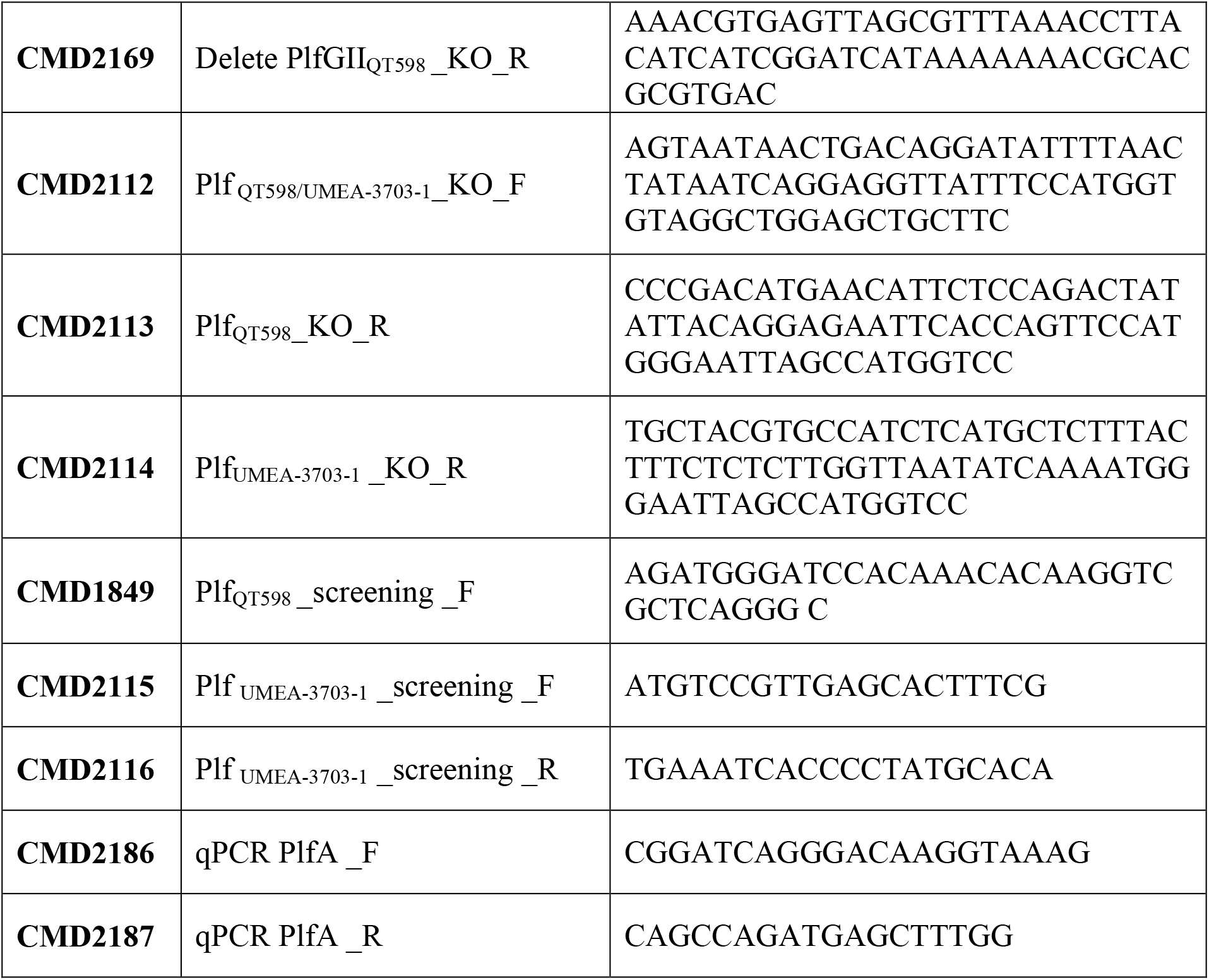
Primers used in this study.

### Deletion of the *plf* genes from strains QT598 and UMEA-3703-1

A *plf* knockout mutant of APEC strain QT598 was obtained by the lamda red recombinase method (Datsenko *et al*., 2000). First, plasmid pKD46 expressing lambda red recombinase was transformed into QT598 by electroporation, then; the kanamycin resistance cassette was amplified from plasmid pKD4 by PCR with primers CMD2112_F and CMD2113_R and transformed into QT598 carrying plasmid pKD46 by electroporation. Mutants were selected at 37°C and then the loss of genes was confirmed by PCR using screening primers CMD1849_F and CMD1900_R, to obtain the QT598Δ*plf* strain (QT4420). The same method was used to create the Δ*plf* deletion mutation in UMEA-3703-1 using primers CMD2112_F and CMD2114_R, to generate strain QT4598 (UMEA-3703-1 Δ*plf*). The deletion was confirmed using primers CMD2115_F and CMD2116_R.

### Extraction of fimbriae and Western blotting analysis

Fimbriae were extracted using the heat extraction method as described previously by (Lymberopoulos *et al*., 2006), with some modifications. Briefly, overnight cultures were incubated at 56°C for 1h and harvested by centrifugation at 4000 rpm for 15 min. Supernatants were incubated in 10% trichloroacetic acid (TCA) to precipitate proteins. Proteins were then concentrated by centrifugation at 12000 rpm for 20 min, washed twice with Tris-EDTA (0.05 M) pH 12 and pH 8.5 and resuspended in 0.1 mL of Tris-EDTA (0.05M) pH 8.5. Western blotting was performed as previously described (Crépin *et al*., 2008) with some modifications. Proteins were separated using 15% polyacrylamide gel, transferred onto a nitrocellulose membrane (Bio-Rad Laboratories, CA, USA), blocked with 15 ml of blocking buffer TBS-T (0,15 M NaCl; 0,025M Tris; 0,05% Tween ; 3% skim milk) for 1h at 4°C. Fimbrial major subunit protein was detected using rabbit polyclonal antibodies provided by New England Peptide (1:1000) against a peptide corresponding to part of the PlfA major fimbrial subunit (Ac-CAHLAADGISVKKD-amide) for 45 min at room temperature. The membrane was then washed 3 times with washing buffer (0,15 M NaCl; 0,025M Tris; 0,05% Tween) and incubated with an anti-rabbit conjugated secondary antibody (1:20000) for 45 min at room temperature. After four washes with TBS-T, proteins were detected using SuperSignal West Pico chemiluminescent substrate (Pierce) according to the manufacturer’s instructions.

### Transmission electron microscopy (TEM)

Bacteria for electron microscopy were grown overnight at 37°C. Cultures were then adsorbed onto a glow-discharged Formvar-coated copper grid for 1 minute and stained with 1% phosphotungstic acid. The excess of liquid was removed with a filter paper. Samples were then dried and observed under a Hitachi H700 transmission electron microscope.

### Hemagglutination assay (HA)

Hemagglutination was performed in 96-well round-bottom plates as described in (Provence *et al*., 1994). Briefly, different types of blood were tested for this assay, human (A and O), horse, bovine, sheep, pig, rabbit, chicken, turkey, and dog red blood cells (RBCs) were suspended in PBS at a final concentration of 3% and added to 96-well plates. Clones expressing different classes of PapG were grown in LB broth at 37°C, centrifuged at 3000 x*g* for 15 min, and pellets were suspended in phosphate-buffered saline (PBS, pH 7.4) and adjusted to an optical density (OD_600_) of 60. The agglutinating titer was determined as the most diluted well with agglutination after 30 min of incubation on ice.

### Biofilm assays

Biofilm formation in 96-well microtiter plates was performed as previously described (Genevaux *et al*., 1996). Fimbrial clones were grown statically in LB at 25°C, 30°C, 37°C and 42°C for 48 hours. After 48h of incubation, the liquid was discarded and plates were washed and stained with 0.1% crystal violet (Sigma) for 15 min. Ethanol-acetone solution (80:20) was used to dissolve biofilm and the optical density was measured at 595 nm to determine the production of biofilm.

### Bacterial adherence to epithelial cell lines

Human bladder 5637 (ATCC HTB-9) and kidney HEK293 (ATCC® CRL-1573™) epithelial cell lines were grown to confluence in 24-well plates in RPMI 1640 or EMEM (Wisent Bio Products, St-Bruno, Canada) supplemented with 10% fetal bovine serum (FBS) at 37°C in 5% CO_2_. Fimbrial clones expressing different classes of PapG or PlfG adhesins were grown in LB medium at 37°C, cultures were then centrifuged, resuspended in RPMI 1640 or EMEM with 10% FBS and added to cells at a multiplicity of infection (MOI) of 10 for 2 h, as described (Matter *et al*., 2011). After 2 h, cells were washed three times with PBS, lysed with 1% Triton X-100, diluted, and plated onto LB agar plates supplemented with selective antibiotics.

### Murine Urinary tract infection model

To determine the potential role of PL fimbriae in virulence, wild type strains QT598 and UMEA-3703-1 as well as the QT598Δ*plf* (QT4420) and UMEA-3703-1Δ*plf* (QT4598) were tested in 6-week-old CBA/J female mice using an ascending UTI model adapted from (Hagberg *et al*., 1983). A total of 5 mice in each group were infected with 10^9^ CFU/ml of bacteria. After 48h, the infected mice were euthanized and bladders and kidneys were harvested aseptically for the bacterial count on MacConkey agar plates. To study the expression of *plf in vivo*, bladder samples after necropsy were homogenized with TRIzol® LS reagent (Thermo Fisher Scientific) for RNA extractions.

### qRT-PCR to measure PL fimbrial gene expression levels

We compared the expression of PL fimbriae by comparing RNA levels of the *plfA* gene in different conditions: LB medium, minimal M63 medium, and during infection in bladders of mice. For *in vitro* analysis, total RNAs from bacterial samples were extracted according to the manufacturer’s protocol EZ-10 Spin Column Total RNA Miniprep Kit (BioBasic). For *in vivo* analysis, bladder samples were homogenized with TRIzol^®^ LS reagent (Thermo Fisher Scientific), incubated with chloroform followed by centrifugation and incubation in ethanol (95-100%) to separate the aqueous phase that contains RNA. Then, RNA samples were extracted using Direct-zol RNA Miniprep kit (Zymo Research, Irvine, CA, USA) according to the manufacturer’s recommendations. All RNA samples were treated with TURBO Dnase (Ambion), to eliminate any DNA contamination. The Iscript^TM^ Reverse transcription supermix (Bio-Rad Life Science, Mississauga, ON, Canada) was used to synthesize cDNAs from samples according to the manufacturer’s protocol. Primers were specific to the *plfA* gene and the RNA polymerase sigma factor *rpoD* (house-keeping control).

qRT-PCR was performed in the Corbett Rotorgene (Thermo Fisher) instrument using 50 ng of cDNA, 100 nM of each primer and 10μl of SsoFast Evagreen supermix (Bio-rad). Data were analyzed using the 2^−ΔΔ*CT*^ (Livak *et al*., 2001).

### Statistical analyses

All data were analyzed with the Graph Pad Prism 6 software (GraphPad Software, San Diego, CA, USA). A Mann-Whitney test was used for mouse infection experiments to determine statistical significance. Analysis of variance (ANOVA) was used to compare the means of samples. Differences between groups were considered significant for P values of p <0.05.

### Ethics statement

Protocols for mice urinary tract infection was approved by the animal ethics evaluation committee – *Comité Institutionel de Protection des Animaux* (CIPA No 1608–02) of the INRS-Centre Armand-Frappier Santé Biotechnologie.

## Acknowledgments

We thank Prof. James Johnson for providing reference clones carrying different classes of Pap and related fimbrial adhesins and and Prof. Niels Frimodt-Møller for providing UPEC strain UMEA-3703-1. We thank Micheline Letarte and Arnaldo Nakamura for assistance with electron microscopy. Funding for this work was supported by Natural Sciences and Engineering Research Council (NSERC) Canada Discovery Grant 2019-06642.

